# Vestibular and auditory hair cell regeneration following targeted ablation of hair cells with diphtheria toxin in zebrafish

**DOI:** 10.1101/2021.06.08.447587

**Authors:** Erin Jimenez, Claire C. Slevin, Luis Colón-Cruz, Shawn M. Burgess

## Abstract

Millions of Americans experience hearing or balance disorders due to loss of hair cells in the inner ear. The hair cells are mechanosensory receptors used in the auditory and vestibular organs of all vertebrates as well as the lateral line systems of aquatic vertebrates. In zebrafish and other non-mammalian vertebrates, hair cells turn over during homeostasis and regenerate completely after being destroyed or damaged by acoustic or chemical exposure. However in mammals, destroying or damaging hair cells results in permanent impairments to hearing or balance. We sought an improved method for studying hair cell damage and regeneration in adult aquatic vertebrates by generating a transgenic zebrafish with the capacity for targeted and inducible hair cell ablation *in vivo*. This model expresses the human diphtheria toxin receptor (hDTR) gene under the control of the *myo6b* promoter, resulting in hDTR expressed only in hair cells. Cell ablation is achieved by an intraperitoneal injection of diphtheria toxin (DT) in adult zebrafish or DT dissolved in the water for larvae. In the lateral line of 5 dpf zebrafish, ablation of hair cells by DT treatment occurred within 2 days in a dose-dependent manner. Similarly, in adult utricles and saccules, a single intraperitoneal injection of 0.05 ng DT caused complete loss of hair cells in the utricle and saccule by 5 days post-injection. Full hair cell regeneration was observed for the lateral line and the inner ear tissues. This study introduces a new method for efficient conditional hair cell ablation in adult zebrafish inner ear sensory epithelia (utricles and saccules) and demonstrates that zebrafish hair cells will regenerate *in vivo* after this treatment.

## Introduction

Loss of hearing or balance can be debilitating and imposes a significant personal, societal, and economic burden upon individuals, their families and communities. Approximately 37.5 million Americans report some degree of hearing loss with incidences increasing with age. 33.4 million adults reported a problem during the past 12 months with balance, unsteadiness, or blurred vision after moving their head. Hearing and balance disorders are most often attributed to loss or damage to the sensory hair cells of the auditory and vestibular organs. The hair cells are mechanosensory receptors that receive signals from our environment and transmit them to the brain. These hair cells reside in the sensory epithelia of auditory and vestibular organs in all vertebrates as well as in the lateral line systems of aquatic vertebrates (Popper and Fay, 1993; Bever and Fekete, 2002; Nicolson, 2005). In zebrafish and non-mammalian vertebrates, hair cells turn over during homeostasis and regenerate completely after being damaged or destroyed by acoustic or chemical exposure, while in mammals, destroying or damaging hair cells results in permanent impairments to hearing and balance. Mammalian hair cell regeneration has been observed but in a very limited fashion in the auditory and vestibular organs of embryonic and newborn mice and mature adults (Burns et al., 2012; Golub et al., 2012; Bucks et al., 2017). Since the majority of hearing and balance disorders in humans are due to the loss or damage of hair cells, understanding how to stimulate the hair cell regeneration process in the mammalian inner ear represents a direct solution to hearing loss or vestibular problems.

The zebrafish is an excellent genetic model to understand hair cell regeneration and inner ear function. Yet, the majority of zebrafish hair cell regeneration and inner ear research to date has focused on the larval lateral line due to its relatively simple structure and accessibility. Lateral line regeneration occurs through support cell proliferation and differentiation. Larval lateral line hair cells frequently undergo apoptosis, constant turnover, and are renewed by peripheral supporting cell division. In the adult inner ear, the sensory epithelium continues to expand for the first 10 months of life and subsequently have a low but measurable turnover of hair cells (Higgs et al., 2002).

To uncover detailed mechanisms of hair cell regeneration in animals that possess the capacity to regenerate hearing, frequently used models for experimental hair cell destruction in adult fish include acoustic overstimulation/sound exposure (Smith et al., 2004; Schuck et al., 2009; Liang et al., 2012), blast wave exposure (Wang et al., 2019), and aminoglycoside antibiotics (Uribe et al., 2013). However, sound exposure experiments achieved ≤75% hair cell ablation and only in the auditory organs while blast wave exposure elicited more serious hearing loss phenotypes, but also caused brain injury with increased cell apoptosis and decreased neurogenesis. Aminoglycoside administration using a high dose of gentamicin induced only a 15% reduction in sensory hair cell loss across the entire saccule and utricle.

In order to investigate the mechanism of hair cell regeneration in adult auditory and vestibular organs, we sought to establish a robust new model for hearing loss and regeneration research by generating a transgenic zebrafish with the capacity for targeted and inducible hair cell ablation in vivo. The gene encoding the human diphtheria toxin receptor was placed downstream of the zebrafish *myo6b* promoter whose expression is limited to differentiated hair cells in zebrafish. Since the orthologous zebrafish receptor has a significantly lower affinity to diphtheria toxin than the human one, treatment with diphtheria toxin results in hair cell specific ablation with minimal systemic side effects. Here we show that a single, low concentration, intraperitoneal injection of DT in the Tg(*myo6b*:hDTR) transgenic background caused complete loss of hair cells in the adult zebrafish utricle and saccule and that over time the hair cells regenerated. We also show that diphtheria toxin exposure ablated larval lateral line hair cells in a dose-dependent manner. This ablation approach could also be used in other tissues where cell-specific ablation is desirable in zebrafish.

## Materials and Methods

### Experimental Animals

TAB5 wild-type (WT) zebrafish were used in this study. Fish were randomly selected and represented roughly equal numbers of males and females. All animal experiments were approved by the Institutional Animal Care and Use Committee under Animal Study Protocol: G-01-3.

### Generation of myo6b-DTR zebrafish

We targeted expression of the human diphtheria toxin receptor (heparin-binding epidermal growth factor precursor; proHB-EGF) to zebrafish hair cells by using the hair cell-specific *myo6b* promoter (Kindt et al., 2012). We generated the Tg(*myo6b*:hDTR) construct as follows. The full coding region of the hDTR gene (Genscript Clone ID OHu26607D) was PCR amplified with the following 5’ adaptor (attB) sequences to increase the specificity of cloning orientation (Forward primer adds attB1 site 5’-GGGGACAAGTTTGTACAAAAAAGCAGGCTTAACCCACTGCTTACTGGCTTA-3’ and Reverse Primer adds attB2 site and a stop codon 5’-GGGGACCACTTTGTACAAGAAAGCTGGGTACTAACTAGAAGGCACAGTCGAGG-3’. A middle entry clone was generated by performing a BP recombination between the attB hDTR PCR product with the pDONR 221 clone. The hDTR-pDONR221 middle entry clone was verified by restriction digest and sequencing. To create a Gateway expression clone, an LR reaction was performed using a 5’ Entry *myo6b* clone (gift from Dr. Katie Kindt), hDTR middle entry clone, 3’ Entry PolyA clone, and destination vector pDestTol2CG2. The *myo6b*:hDTR construct was verified by sequencing and injected into TAB5 1-cell stage embryos with mRNA encoding the Tol2 transposase (Kawakami et al., 2000). All experimental zebrafish used in this study were heterozygous, Tg(*myo6b*:hDTR).

### In-Situ Hybridization Chain Reaction (HCR)

3 dpf embryos were fixed in a 4% formaldehyde solution and stored overnight in 100% methanol. Embryos were rehydrated with a series of graded 1 mL methanol/PBST washes (75% methanol, 50% methanol, 25% methanol) for 5 minutes and then rinsed 5 times in PBST (0.1% Tween-20) for 5 min. Embryos were treated with 1 mL of proteinase K (2 mg/mL) for 10 minutes at room temperature followed by two washes with PBST without incubation. Embryos were then postfixed with a 4% formaldehyde solution for 20 minutes at room temperature. Following fixation, embryos were washed 5 times for 5 minutes with PBST. Embryos were hybridized with HCR probes that were purchased commercially (Molecular Instruments, Inc.) and targeted hDTR (HBEGF, NM_001945.3) and Myo6b (NM_001004110.1). Detection and amplification was performed in accordance with the HCR v3.0 protocol for whole-mount zebrafish embryos and larvae (Choi et al., 2016). Embryos were mounted in 0.8% low melting agarose and imaged on a Zeiss LSM 880 confocal microscope. Maximum intensity projections of z-stacks were generated in Image J/Fiji software (Schindelin et al., 2012).

### RNA isolation from adult inner ear tissues

Saccules and utricles were dissected from adult wild-type and Tg(*myo6b*:hDTR) zebrafish and homogenized in 0.7 mL TRIzol Reagent (Thermo Fisher Scientific, USA) with a power homogenizer. RNA was isolated from aqueous phase after TRIzol/chloroform extraction and treated with DNase I. RNA was purified using the RNA Clean & Concentrator-5 (Zymo Research) and measured (Nanodrop One).

### Quantitative real-time PCR analysis (RT-qPCR) on adult inner ear tissues

RNA was transcribed into cDNA according to manufacturer’s instructions (SuperScript III RT, Thermo Fisher Scientific, USA). RT-qPCR was performed in technical replicates using 1:1 cDNA in each reaction and a primer concentration of 0.5uM. PowerUp SYBR Green Master Mix (Thermo Fisher Scientific Cat, #4344463, USA) and self-designed primers were used (Eurofins, Luxembourg). Primers were designed by using Primer3 followed by a UCSC In-Silico PCR to search the zebrafish sequence database. hDTR was amplified using the forward primer 5’ - GACCCTCCCACTGTATCCAC - 3’ and the reverse primer 5’ - GCTCCTCCTTGTTTGGTGTG - 3’. myo6b was amplified using the forward primer 5’ - ATTAAGAGCTATCAGGGACGC - 3’ and the reverse primer 5’ - GCTCATCTTCAGAACCCTCAT - 3’. ef1alpha was used as a housekeeping gene and was amplified using the forward primer 5’ - CGACAAGAGAACCATCGAGAAGTT - 3’ and the reverse primer 5’ - CCAGGCGTACTTGAAGGA - 3’.

### Larval Zebrafish Diphtheria Toxin Treatments

Diphtheria toxin was purchased from Sigma-Aldrich (D0564). 5 dpf transgenic larvae were exposed for 3 h to 12 h in various concentrations of diphtheria toxin dissolved in 1X Holtfreter’s medium, washed and then maintained in 1X Holtfreter’s for up to 3 days. 8 dpf zebrafish were treated with 8 μM YO-PRO-1 dye (Y3603, Molecular Probes, OR, USA) dissolved in 1X Holtfreter’s medium for 1 hour at 28.5°C. After washing, fish were lightly anaesthetized with 0.01% tricaine and placed in 96-well, glass bottomed plates for observation. Stained neuromasts in the lateral trunk region were visualized and quantified with an inverted Leica stereomicroscope using a 10X objective. For time-course experiments, larvae were returned to 1X Holtfreter’s medium for recovery.

### Adult Zebrafish Diphtheria Toxin Administration

Diphtheria toxin (DT) was dissolved in 1XPBS. 6 to 10 month-old wild-type (TAB5) and transgenic adult zebrafish of mixed sex were injected one time with diphtheria toxin into the abdominal cavity, posterior to the pelvic girdle, using a microsyringe for nanoliter injection with a 35G beveled needle. Concentration ranges from 0.01 ng to 50 ng per fish were tested. Before intraperitoneal injection, fish were fasted for 24 hours and then lightly anesthetized with buffered MS-222. Immediately after injection, fish recovered in fresh system water and maintained off system for up to 14 days. Fish were fed and water changed once daily. Health and water quality inspections were completed twice daily.

### Histological Methods

Adult zebrafish were euthanized using buffered MS-222. The heads were dissected and fixed in 4% formaldehyde overnight at 4°C. Inner ears were dissected as previously described in Liang and Burgess, 2009.

### Hair Cell Labelling

Alexa Fluor 488 phalloidin was used to visualize and quantify F-actin in stereocilia of zebrafish. Utricles and saccules were dissected and stained using Alexa Fluor 488 phalloidin as previously described in Liang, 2009 and 2012. Proteins were detected in whole-mount utricles and saccules using standard immunofluorescence labeling methods. Following overnight fixation, inner ear sensory epithelia were rinsed several times in PBTX (PBS plus 0.1% Triton X-100) and blocked for 1 hour in BBTX at room temperature (PBS plus 0.5% BSA, 2% NGS, and 0.1% Triton X-100). Inner ear sensory epithelia were incubated overnight at 4°C with primary antibodies. Inner ears were washed three times for 10 minutes in PBTX and then incubated overnight at 4°C with secondary antibodies in BBTX. After washing three times for 10 minutes in PBTX, inner ear sensory epithelia were incubated for 45 minutes at room temperature with Alexa Flour 488 phalloidin in PBS (Thermo Fisher Scientific, #A12379, USA, 1:1000-dilution). Following three washes in PBS for 10 minutes, saccules (with or without lagena) and utricles were mounted in Vectashield with DAPI. Primary and secondary antibodies used include the rabbit myosin VI and myosin VIIa antibodies (Proteus Biosciences 25-6790 & 25-6791, 1:300-dilution), rabbit cleaved Caspase-3 (Cell Signaling #9661, 1:300-dilution), Alexa Fluor 568 goat anti-rabbit IgG (1:1000-dilution).

### Cellular Imaging and Analysis

Confocal images were acquired with a Zeiss LSM 880 confocal microscope. Confocal Z stacks of the entire saccule and utricle were projected into a single image to capture all phalloidin positive cells from different planes of focus for hair cell counting. Counts of phalloidin labelled hair cell bundles were obtained from preselected 50 μmx50 μm digital boxes along the rostral-caudal axis of the saccule or the medial-striola planes of the utricle. Normal hair cells were quantified as hair cell bundles with intact stereocilia using Image J/Fiji software (Schindelin et al., 2012). Cleaved caspase-3 positive cells were counted with Image J/Fiji software from the entire saccule and utricle whole mount.

## Results

### Generation of myo6b:hDTR zebrafish

To create a model that would allow for hair cell specific ablation, we utilized the human DTR gene, which has previously been used to effectively ablate hair cells in the mouse utricle (Golub et al., 2012). In order to drive expression of hDTR only in hair cells, we use the zebrafish hair cell specific promoter *myo6b* which is expressed in auditory, vestibular, and lateral line hair cells in zebrafish (**Figure 1A**) (Obholzer et al., 2008, Matern et al., 2018). The construct was cloned using the Gateway System into a Tol2 transposon vector and was injected with Tol2 transposase mRNA (Kawakami et al., 2000) into zebrafish embryos at the one-cell stage to create a stable transgenic line. To confirm Tg(*myo6b*:hDTR) was expressed in hair cells, we performed *in situ* hybridization chain reaction (HCR) on wild-type and Tg(*myo6b*:hDTR) 3 dpf embryos using a probe targeting the human diphtheria toxin receptor (hDTR). As expected, the hDTR fluorescent signal was present in lateral line hair cells and in the anterior macula of Tg(*myo6b*:hDTR) fish, but absent in wild-type controls. As a control, we simultaneously hybridized and detected a probe for myo6b in lateral line hair cells and in the anterior macula of the inner ear of Tg(*myo6b*:hDTR) and wild-type fish (**Figure 1C,D**). hDTR expression was also verified by qRT-PCR in adult zebrafish inner ear auditory (saccule) and vestibular (utricle) sensory epithelia of stable transgenic lines (**Figure 1B**).

**Figure 1.**
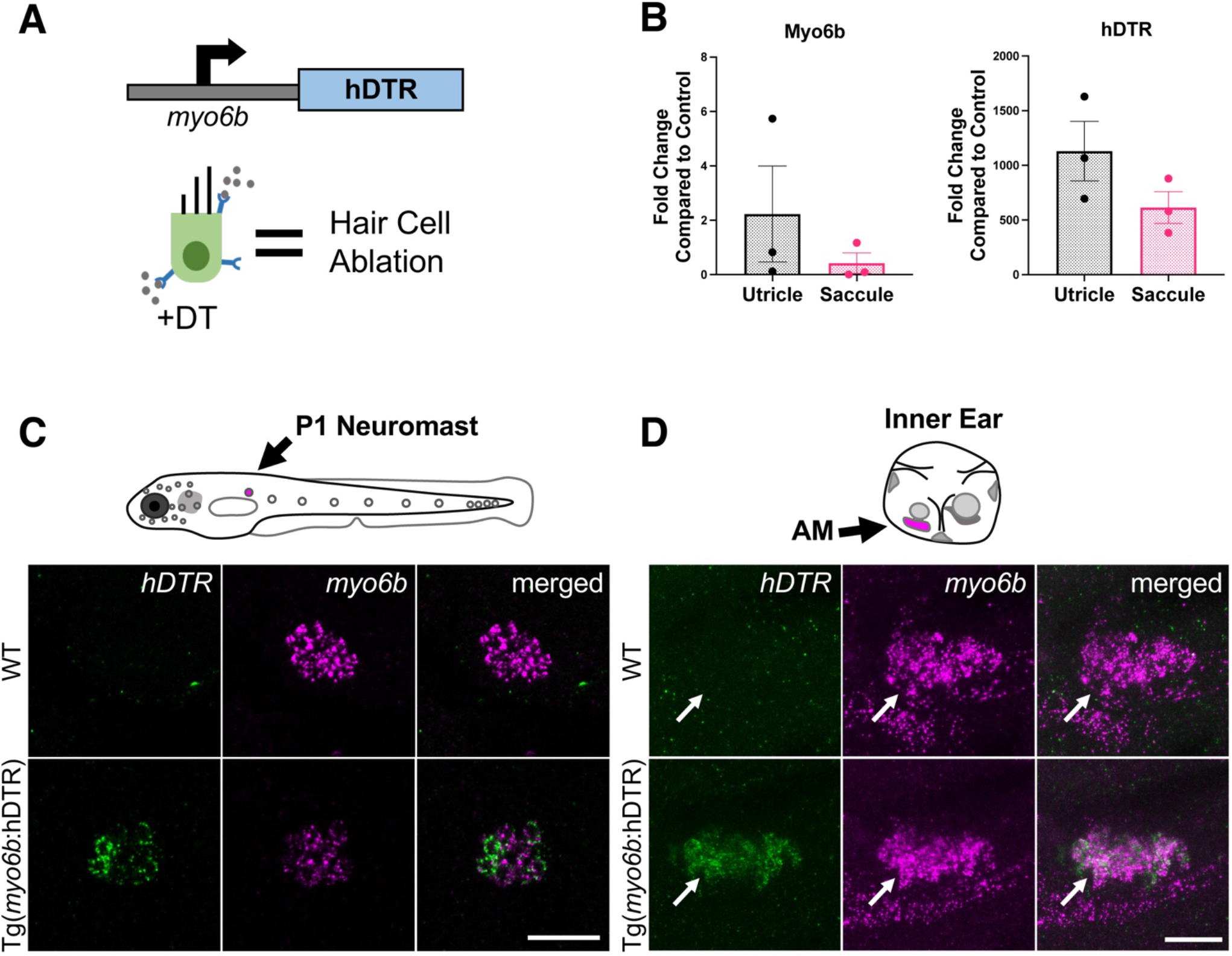
The Tg(*myo6b*:hDTR) zebrafish. **(A)** Schematic representation of the hDTR construct driven by the hair cell-specific *myo6b* promoter and representation of the hair cell-specific ablation approach. DT is internalized in cells specifically expressing the human version of the diphtheria toxin receptor, triggering cell death. **(B)** qRT-PCR analysis on untreated Tg(*myo6b*:hDTR) zebrafish saccule and utricle showing expression of *hDTR* and *myo6b* relative to wild-type animals. The values are represented as the mean ± SEM from 3 independent samples. **(C)** Schematic depicts a lateral view of a whole zebrafish larva with neuromasts (gray circles). The P1 neuromast examined for imaging is indicated by magenta fill. *In situ* HCR using probes targeting *hDTR* and *myo6b* in hair cells of a single P1 neuromast of 3 dpf wild-type (WT) and Tg(*myo6b*:hDTR) embryos is shown. Brightness and contrast adjusted 50% and 25%, respectively. Scale bar 20 μm. **(D)** Schematic depicts the left ear of a larval zebrafish with anterior macula indicated by magenta fill. Lateral views of inner ears are shown following HCR *in situ* with probes targeting *hDTR* and *myo6b* in 3 dpf wild-type (WT) and Tg(*myo6b*:hDTR) embryos. Brightness and contrast adjusted 65% and 35%, respectively. Scale bar 20 μm.

### DT induced hair cell death and regeneration in the larval lateral line

To determine whether DT was able to ablate hair cells of the lateral line of zebrafish expressing hDTR in hair cells, we performed a dose-response assay using 5 dpf larvae that were wild-type or heterozygous for the Tg(*myo6b*:hDTR) allele. We treated 5 dpf larvae with concentrations of dissolved DT ranging from 0.5 μg/mL to 1.5 μg/mL and continuous exposure times ranging from 3 h to 12 h (**Figure 2C**). To examine the dose-response relationship, larval neuromasts were labeled with YO-PRO-1 before and after exposure to DT. The number of YO-PRO-1 labeled cells in each of the four identified neuromasts (P1, P2, P3, P4) was determined for the different exposure concentrations and durations (**Figure 2C**). Hair cells were assessed by fluorescence microscopy and counted for approximately 8 fish per group.

**Figure 2.**
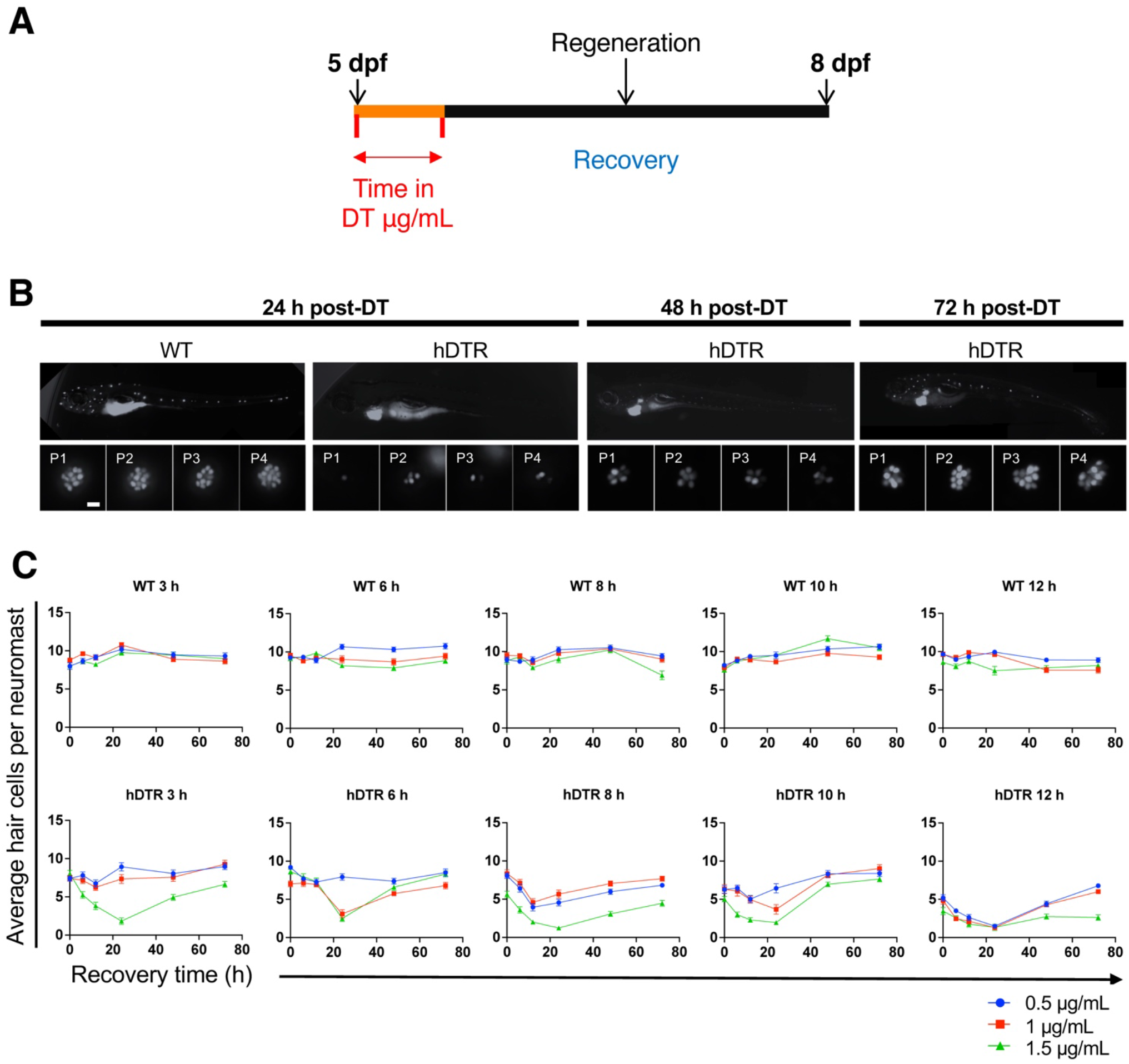
Tg(*myo6b*:hDTR) larval zebrafish show hair cell loss and regeneration in lateral line neuromasts after *in vivo* DT treatment. **(A)** Schematic representation of the larval DT exposure treatment. **(B)** Regeneration time-course showing P1-P4 over three days of recovery. Larvae were exposed to 1 μg/mL of DT for 6 h. Scale bar 100 μm. **(C)** Wild-type (WT) and Tg(*myo6b*:hDTR) (hDTR) larvae were exposed to three concentrations of DT (0.5 μg/mL, 1 μg/mL, and 1.5 μg/mL) for various durations (3 h, 6 h, 8 h, 10 h, 12 h). Neuromast viability was monitored daily by YO-PRO-1 labelling 0 h to 72 h post-incubation. Shown is the quantification of YO-PRO-1 labeled hair cells. The values are represented as the mean ± SEM from 8 fish.

DT had no effect on YO-PRO-1 staining in wild-type 5 dpf zebrafish (**Figure 2B,C**). However, DT reduced YO-PRO-1 staining in 5 dpf Tg(*myo6b*:hDTR) fish in a dose-dependent manner with no general toxicity or mortality at any of the doses or exposure times (**Figure 2B,C**). A representative neuromast from unexposed zebrafish larvae at 5 dpf and the same neuromast from larvae exposed to 1 μg/mL of DT for 6 h are shown (**Supplementary Figure 1C**). Individual hair cells were brightly stained with YO-PRO-1 in wild-type fish (**Figure 2B, Supplementary Figure 1B**). By comparison, larvae exposed to DT showed reductions in both the intensity of YO-PRO-1 and the number of labeled hair cells. At the higher exposure concentrations and longer exposure times, the YO-PRO-1 staining in the neuromasts was absent **(Figure 2B,C)**.

We observe a delay and cumulative effect of DT on hair cell death which is likely a result of diphtheria toxin’s mechanism of action. The dose and time of exposure influence the amount of hair cell ablation, but synchronous hair cell loss in Tg(*myo6b*:hDTR) continued even after withdrawal of DT. Lower doses and longer exposure times were as effective at ablating hair cells as high doses for shorter times. At 3 h incubation with 1.5 μg/mL of DT, hair cells are progressively lost until 24 h post treatment, but no significant hair cell loss is observed immediately with treatment at 0 h (**Figure 2C**). These data show a relationship between DT concentration, exposure time, and hair cell death.

Lateral line hair cells regenerate following acute chemical injury. In Tg(*myo6b*:hDTR) larvae, hair cells regenerate (defined as recovery in YO-PRO-1 labeling) when DT exposed animals are removed from the toxin. To determine the time-course for recovery, hair cell regeneration in the four identified neuromasts was monitored at 0, 6, 12, 24, 48, and 72 h after exposure to DT. Hair cell regeneration was evident within 2 days of removal (**Figure 2B,C** and **Supplementary Figure 1C)**. After 3 days, the mean number of hair cells per neuromast in the transferred larvae was not significantly different from the unexposed larvae indicating that recovery was complete. In contrast, larvae exposed to DT at the highest (1.5 μg/mL) and longest duration (12 h) showed only partial hair cell regeneration after 3 days.

### DT induced hair cell death and regeneration in the adult inner ear

Using fluorescently-tagged phalloidin to visualize hair cell bundles and myosin VI/VIIa antibodies for hair cell bodies, we assessed utricle and saccule hair cells in untreated adult zebrafish that were wild-type or heterozygous for the Tg(*myo6b*:hDTR) allele (**Supplementary Figure 2**). Fluorescent phalloidin, a highly specific F-actin stain, was used to visualize hair cells. Hair cell bodies were labeled with myosin VI/VIIa (**Supplementary Figure 2**, red channel), which labels the cytoplasm of hair cells to confirm cell death (as opposed to hair cell bundle damage). Figure 3 illustrates the overall appearance of a phalloidin stained utricle and saccule (**Figure 3B**). The average hair cell densities of the untreated Tg(*myo6b*:hDTR) utricle and saccule maculae are 85 and 161 per 2.5 mm^2^ area, respectively. Myosin VI/VIIa was present in the cytoplasm of all hair cells examined in untreated wild-type, DT treated wild-type, and untreated DTR fish (**Supplementary Figure 2**, red channel). The pattern of phalloidin and myosin VI/VIIa labelling was consistent in treated and untreated wild-type controls and untreated Tg(*myo6b*:hDTR) sensory epithelia, although there was some variability in fluorescence intensity that was not quantified (**Figure 4, Supplementary Figure 2**) (Coffin, et al., 2007). The number of hair cells per utricle and saccule did not differ significantly between these two groups and no differences were observed with respect to hair cell appearance. These observations suggested that inner ear hair cell development and maintenance were not affected in Tg(*myo6b*:hDTR) zebrafish.

**Figure 3.**
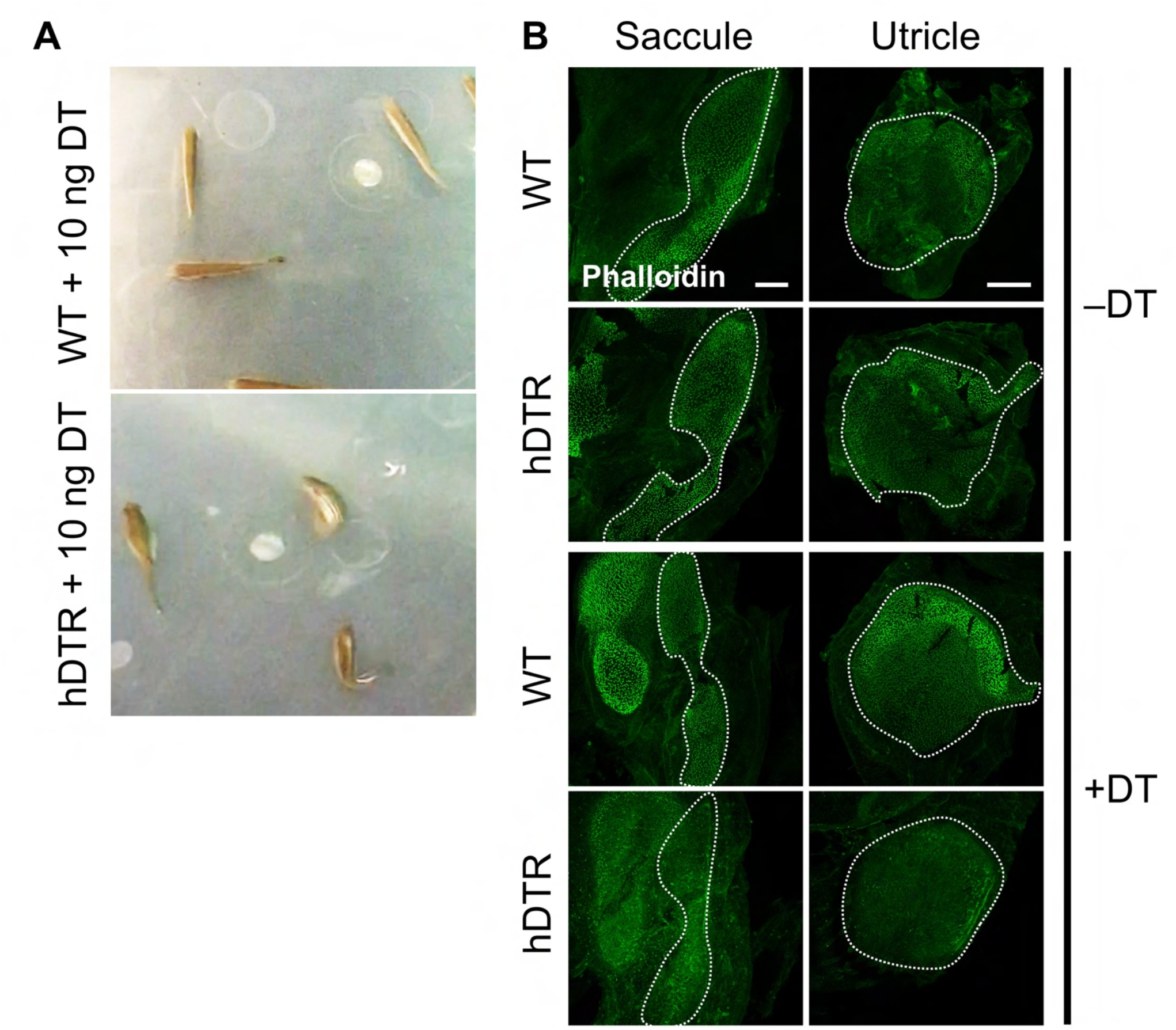
Adult Tg(*myo6b*:hDTR) injected fish exhibit spatial disorientation and impaired balance. **(A)** Injected wild-type fish (WT, top) and Tg(*myo6b*:hDTR) (hDTR, bottom) with 10 ng of DT 3 days-post injection. **(B)** Saccule and utricle from untreated (-DT) or treated (+DT) wild-type and Tg(*myo6b*:hDTR) fish. Saccule and utricle from treated(+DT) wild-type and Tg(*myo6b*:hDTR) fish were isolated 3 days post-DT. Scale bar 100 μm.

**Figure 4.**
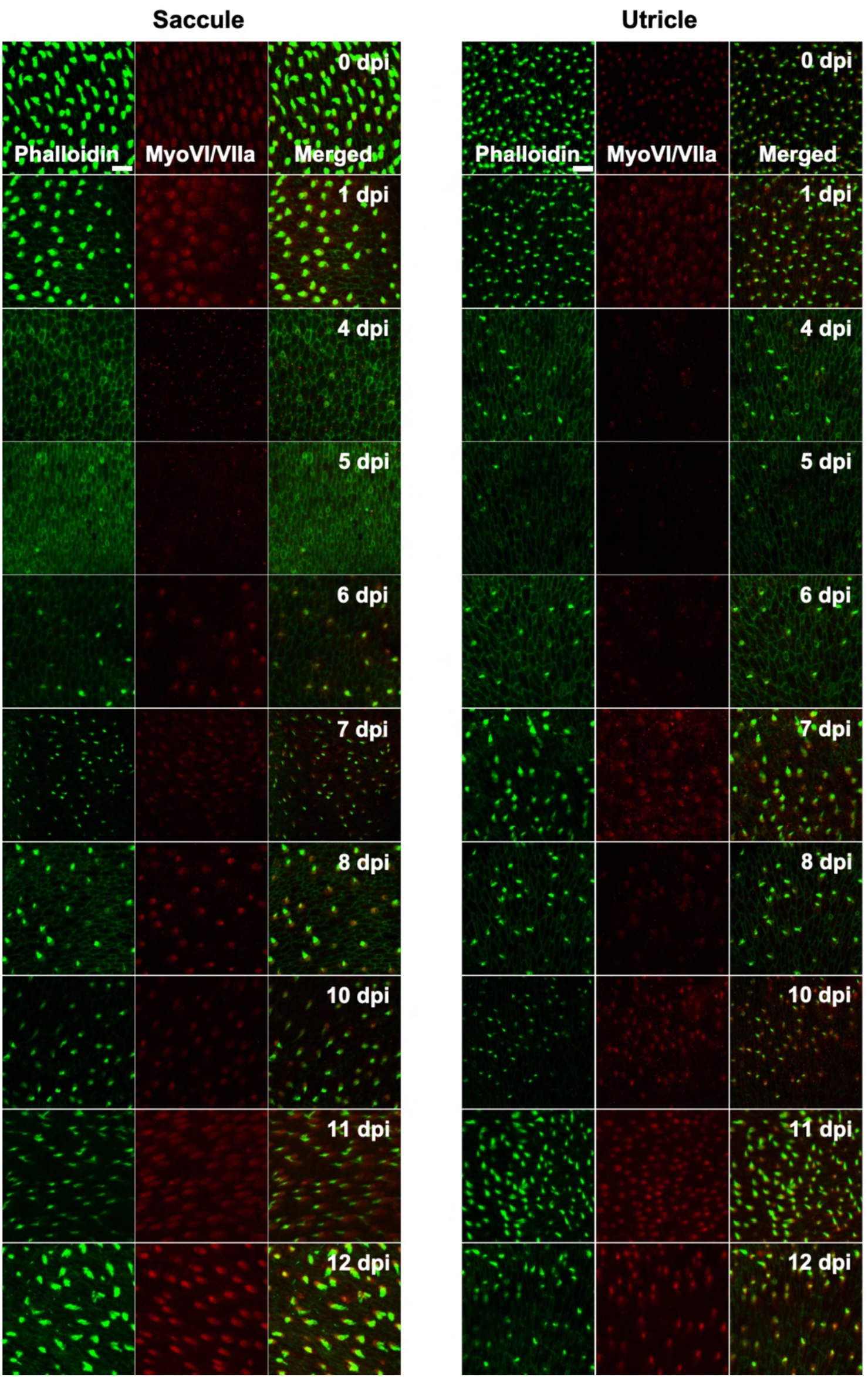
Tg(*myo6b*:hDTR) adult zebrafish hair cell loss and regeneration in the inner ear after DT treatment. Saccule and utricle were isolated at specified timepoints following DT administration in Tg(*myo6b*:hDTR) fish. Close examination of hair cells with phalloidin (green channel) and hair cell bodies with anti-myosin VI/VIIa (red channel). Scale bar 100 μm. For qualitative purposes, brightness was increased 10% and contrast by 20% across all images.

In order to determine an appropriate dose of DT to use for downstream ablation experiments in adult zebrafish, we evaluated the effect of various concentrations of injected DT (ranging from 0.01 ng to 10 ng per fish) on adult fish that were wild-type or heterozygous for the Tg(*myo6b*:hDTR) allele. No to minimal defects in swimming behaviors were observed in Tg(*myo6b*:hDTR) fish injected with 0.01-0.05 ng of DT. At concentrations above 1 ng per fish (such as 10 ng), Tg(*myo6b*:hDTR) fish exhibited consistent swimming defects that come out most strongly when water is agitated (**Supplemental Video 1**). The behavior of injected Tg(*myo6b*:hDTR) fish with 10 ng of DT consisted of somersaulting and random lateral looping (**Figure 3A** of image of dorsal view of swimming patterns of adult wild-type and Tg(*myo6b*:hDTR) injected fish). Tg(*myo6b*:hDTR) injected fish mimic the swimming behavior of fish under microgravity conditions (Von Baumgarten et al., 1975), fish that have undergone laceration or removal of the utricle (Pfeiffer, W., 1964), and the *sputnik* and *orbiter* adult circler mutants with mutations affecting vestibular function (Nicolson et al. 1998). The onset of swimming defects correlated to when the hair cells in the inner ear were maximally ablated (**Figure 3**). Similarly, recovery in swimming behavior correlated to when the hair cells in the inner ear regenerated.

We characterized the time-course of hair cell loss and the minimum dose capable of completely ablating hair cells to help maximize animal viability and reduce excess stress. Similar to our observations in larval zebrafish treated with DT, we observed a delay and cumulative effect of DT on hair cell death in adult zebrafish. The dose of exposure influenced when hair cell loss was first observed and the time it took for hair cells to return. We determined a low DT dose of 0.05 ng per fish was sufficient for complete hair cell ablation throughout the utricle, saccule and even lagena followed by hair cell recovery within a 13-day window.

Adult fish received a single intraperitoneal injection of 1 μl of 0.05 ng DT. Immediately following injection, fish were allowed to recover for up to 14 days in static tanks. Adult inner ears (utricle and saccule) were harvested following timepoints ranging from 0 to 13 days post injection. DT did not affect hair cells in injected wild-type zebrafish at all time points examined. In contrast, significant hair cell loss was observed between 4 and 5 days after DT treatment in adult Tg(*myo6b*:hDTR) zebrafish (**Figure 4,5**). We confirmed elimination of hair cells (as opposed to hair cell bundle damage) by co-labeling sensory epithelia with phalloidin and myosin VI/VIIa antibodies. Phalloidin staining revealed structural changes that occurred at the epithelial surface 4 and 5 days after DT treatment such as putative lesions and bundle-less cuticular plates in the region of stereociliary loss over the time course of hair cell death. The lack of phalloidin in stereocilia and bundle-less cuticular plates correlated with the absence of myosin VI/VIIa labeling (**Figure 4**). We conclude that complete hair cell loss is achieved 5 days post-DT treatment in adult Tg(*myo6b*:hDTR) zebrafish.

**Figure 5.**
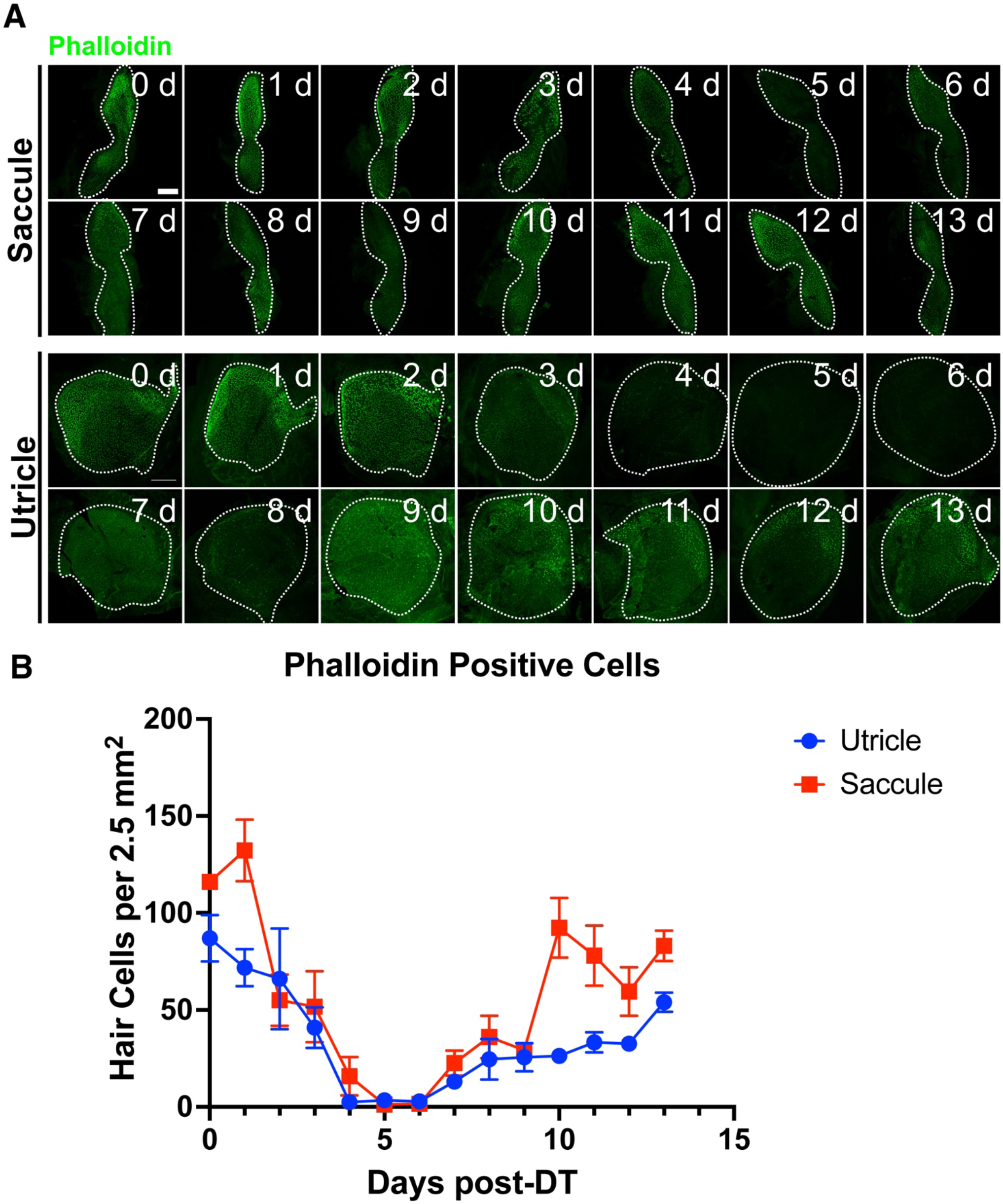
Tg(*myo6b*:hDTR) adult zebrafish hair cell loss and regeneration in the inner ear after DT treatment. **(A)** Tg(*myo6b*:hDTR) saccule and utricle used for quantification were isolated at specified timepoints following DT administration and hair cells were labeled with phalloidin (green channel). Scale bar 100 μm. **(B)** Quantification of phalloidin positive hair cell number after DT injection. Error bars demonstrate the mean ± SEM.

Hair cells re-emerged as indicated by the appearance of short phalloidin positive hair cell bundles and myosin VI/VIIa positive cell bodies on day 6 post-DT treatment. On day 7 post-DT, saccules and utricles exhibited an increase in short, immature-like bundles, and an increase in hair cell bundle density. The increase in phalloidin labelled stereocilia corresponded to an increase in myosin VI/VIIa positive cell bodies (**Figure 4**). The pattern of hair cell bundle density and intensity using phalloidin labeling remained consistent after day 8 in the saccule and utricle, although there was some variability in fluorescence intensity using myosin VI/VIIa labeling. The hair cells return to control levels by 13 days post-DT (**Figure 5**).

### Hair cell apoptosis is triggered by DT treatment in hDTR transgenic fish

Apoptosis results in cleavage of caspase-3. Therefore, to test whether hair cells in treated Tg(*myo6b*:hDTR) fish were undergoing apoptosis, we stained for cleaved caspase-3 on DT treated fish. In the utricle and saccule of DT treated Tg(*myo6b*:hDTR) fish, we observed cleaved caspase-3-positive cells 1 day post-DT and the number of cleaved caspase-3 positive cells significantly increased by day 2 post-DT (**Figure 6**). The number of cleaved caspase-3 positive cells declined on day 3 post-DT and each day thereafter examined (**Figure 6B**). We conclude that at least some of the hair cell death is due to apoptosis and cell death is initiated as early as 24 hours post DT treatment.

**Figure 6.**
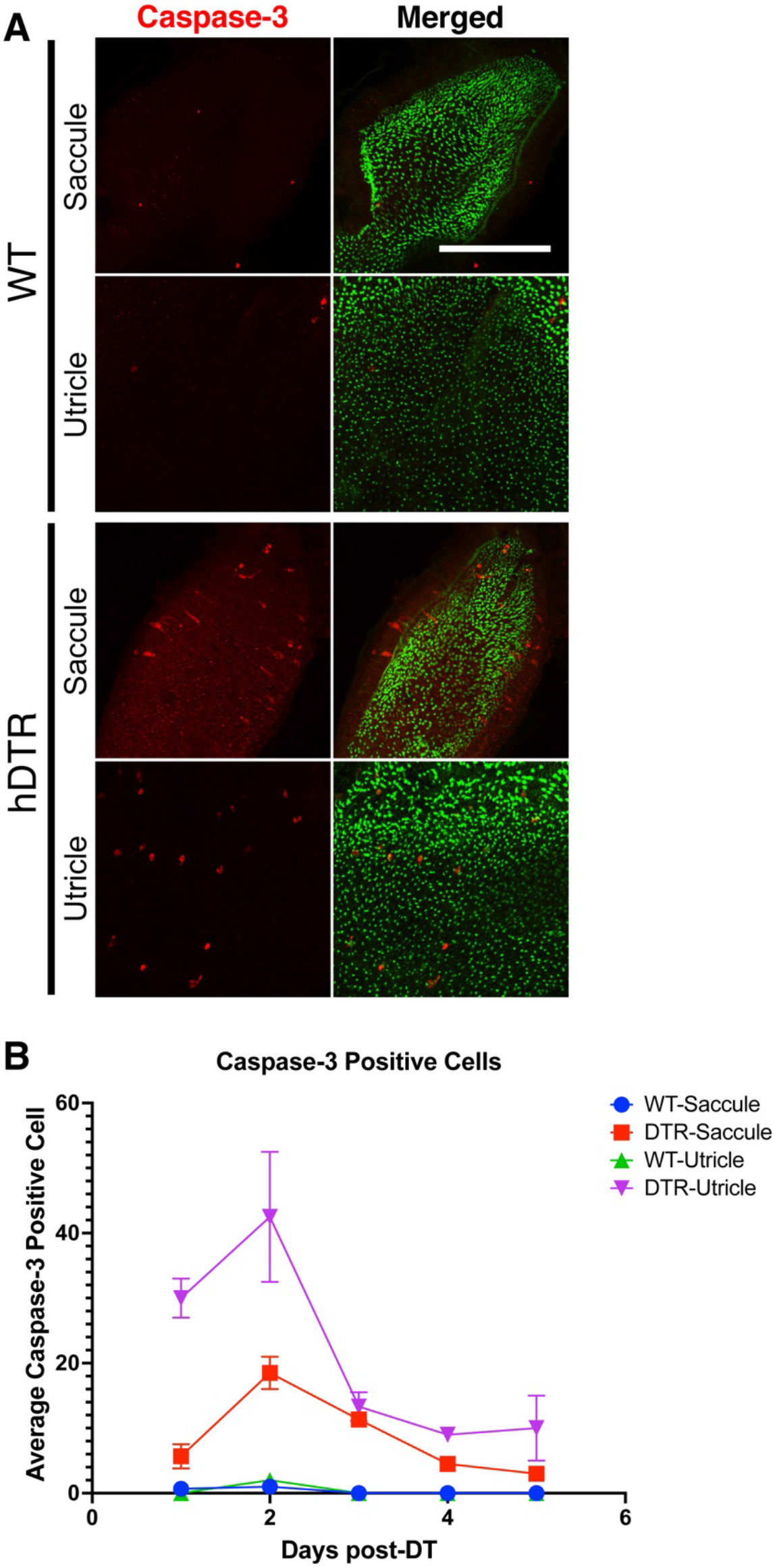
DT induced hair cell death is primarily due to apoptosis. **(A)** Saccule and utricle from wild-type (WT) and Tg(*myo6b*:hDTR) (hDTR) inner ear tissues 2 days post-DT. Hair cells are labeled with phalloidin (green channel) and cleaved caspase-3 (red channel) positive cells were detected 2 days post-DT. **(B)** Quantification of cleaved caspase-3 positive cells in saccule and utricle dissected on days 1 to 5 following DT injection. Scale bar 100 μm. For qualitative purposes, brightness and contrast increased by 20%.

## Conclusions

We show that a stable zebrafish line expressing the human diphtheria toxin receptor in a tissue-specific fashion, can be successfully applied to the analysis of hair cell regeneration in the zebrafish adult inner ear and larval lateral line systems. Incubation of larvae or adult injection of DT was utilized to achieve hair cell specific ablation in zebrafish expressing hDTR under the control of the *myo6b* promoter. With this technique, we observe a delay in cell death which is likely due to the mechanism of action for diphtheria toxin. DT binds to the toxin receptor, it internalizes, and the catalytic A subunit acts as an inhibitor of protein synthesis and arrests RNA synthesis, which ultimately leads to apoptosis or cell death (Collier R. J., 1975). A potential drawback of the system is the extreme potency of the toxin which can cause “off site injuries.” Studies which express diphtheria toxin A fragment (DTA) in zebrafish embryos in the retina (Kurita et al., 2003) and in germline (Slanchev et al., 2005) resulted in undesired ablation of other cells and even death of the organism due to DT toxicity. However, we find that cell specific expression of the human diphtheria toxin receptor in hair cells using the *myo6b* promoter causes no morphological defects in comparison to wild-type fish. Moreover, diphtheria toxin can be administered at low enough doses to cause cell specific ablation with limited to no toxicity in other cells and no noticeable toxicity to the animal.

### Comparison with other adult zebrafish models for hair cell ablation

At least 3 additional zebrafish models have been used to ablate hair cells in the adult inner ear. Immediately following cessation of sound exposure (to 100 Hz pure tone at 179 dB re 1 μPa RMS) for 36 h results in a 75% reduction in stereocilia density of the auditory hair cells in saccules. These sound exposure experiments significantly decrease hair cells to 43% in the caudal region and 75% of the total distance from the rostral tip. Specifically, sound exposure produces hair bundle loss only in the caudal region of the saccule and then 2 days later noticeable bundle loss is seen in the central portion of the rostral region (25%) 0 and 2 days post sound exposure (Schuck et al., 2009). Similar to our observations using DTR-fish treated with DT, auditory hair cell regeneration is observed 2 days post sound exposure. Aminoglycoside administration using a single intraperitoneal injection of a high dose of gentamicin induces only a noticeable reduction in sensory hair cell loss across the entire saccule and utricle, accompanied by shifts in auditory thresholds (Uribe et al., 2013). It is not shown whether hair cells replenish following gentamicin exposure. In contrast to both acoustic and ototoxic exposure, we observe near complete hair cell loss throughout the auditory and vestibular sensory epithelia followed by regeneration of hair cells. The Tg(*myo6b*:hDTR) zebrafish and DT system represents an ideal method for hair cell ablation in a regeneration study as the method induces synchronous destruction of all hair cells with negligible effects on neighboring inner ear cells. Moreover, this system is reversible to permit hair cell regeneration.

Most hair cell regeneration studies have been implemented using the larval lateral line systems. However, the larval lateral line system and inner ear hair cells are not identical and examination of adult inner ears could have significant differences relevant to potential therapeutic treatments. The Tg(*myo6b*:hDTR) fish will enable examination of adult behaviors associated with auditory dysfunction and equilibrium orientation defects. DT treated Tg(*myo6b*:hDTR) adult zebrafish have defects similar to those seen in human hereditary and environmentally induced forms of deafness which may serve as a model for such disorders since zebrafish are accessible to a wide range and levels of analyses. This method can uncover roles of specific tissues during development, homeostasis, and aging. Analysis of recovery after cell ablation may also reveal novel cellular and molecular mechanisms underlying the regenerative processes, thus bringing insights to the field of regenerative medicine.

## Author Contributions

EJ and SMB designed the experiments and wrote the manuscript. EJ, CS, and LC-C performed the experiments.

## Acknowledgments

We thank Dr. Katie Kindt for generously donating the 5’ entry *myo6b* clone; Blake Carrington and Raman Sood for assistance with cell injections; Stephen Frederickson and Dr. Tannia Clark, and Charles River for zebrafish care; and the members of the Burgess laboratory for helpful discussion. All animal experiments were approved by the National Human Genome Research Institute’s Animal Care and Use Committee (protocol #G-01-3).

## Supplementary Material

### Supplementary Data

**Supplementary Video 1. DT induced hair cell death affects adult swimming behaviors.** Injected wild-type fish (left panel) and Tg(*myo6b*:hDTR) with 10 ng of DT 3 days-post injection. Tg(*myo6b*:hDTR) display summersaulting and lateral looping when water is agitated.

### Supplementary Figures

**Supplementary Figure 1.**
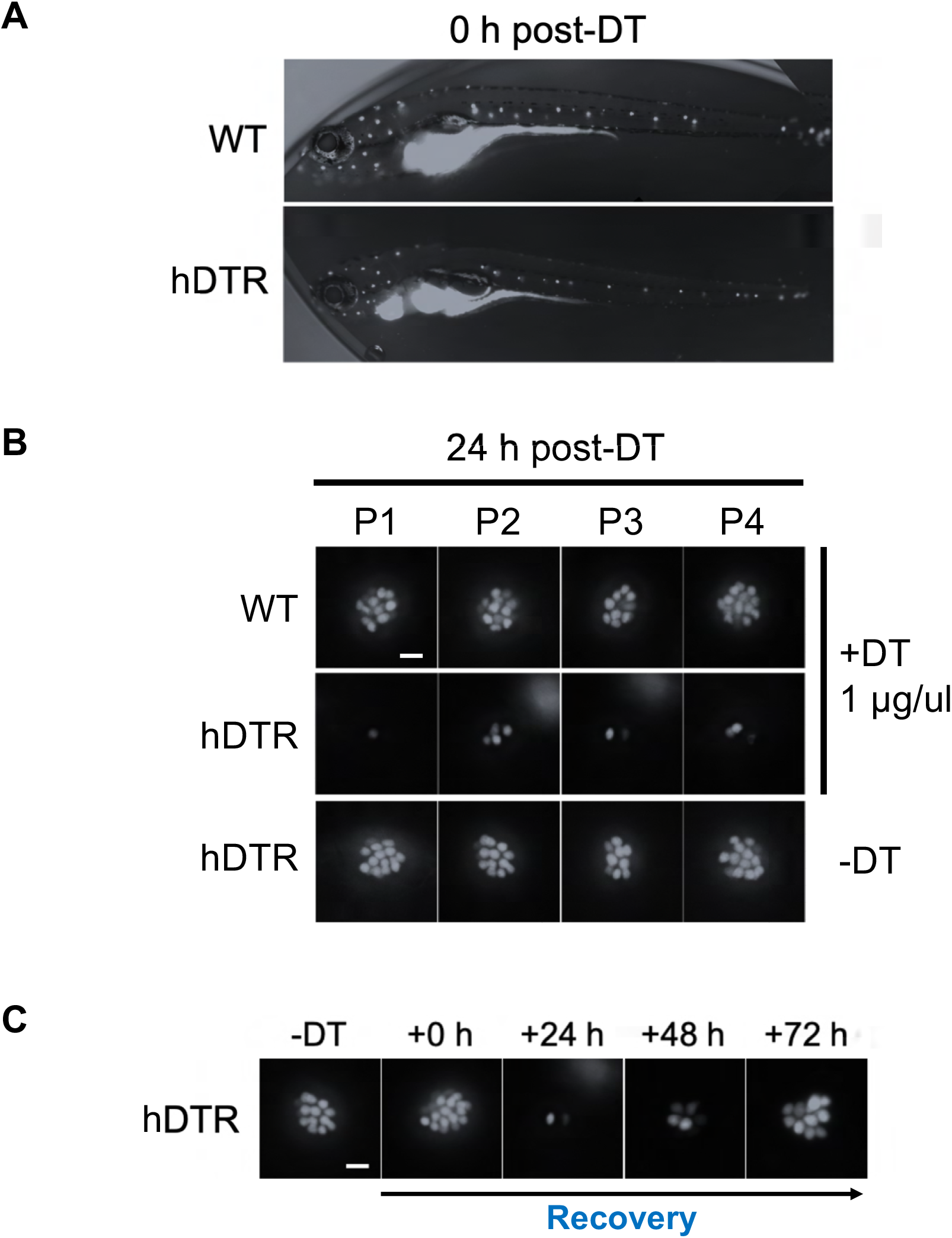
Tg(*myo6b*:hDTR) larval zebrafish show hair cell loss and regeneration in lateral line neuromasts after *in vivo* DT treatment. Larvae were exposed to 1 μg/mL of DT for 6 h. Neuromast viability was monitored daily by YO-PRO-1 labelling 0 h to 72 h post-DT incubation. The 1 μg/mL concentration and the 6 h duration of DT exposure was selected based on dose-dependent work in Figure 2. **(A**) Wild-type (WT) controls and Tg(*myo6b*:hDTR) larvae (hDTR) 0 h post-DT. **(B)** Neuromasts 24 h post-DT in WT and Tg(*myo6b*:hDTR) larvae. **(C)** Regeneration time-course showing P3 neuromast of Tg(*myo6b*:hDTR) larvae treated with DT. Scale bar 100 μm.

**Supplementary Figure 2.**
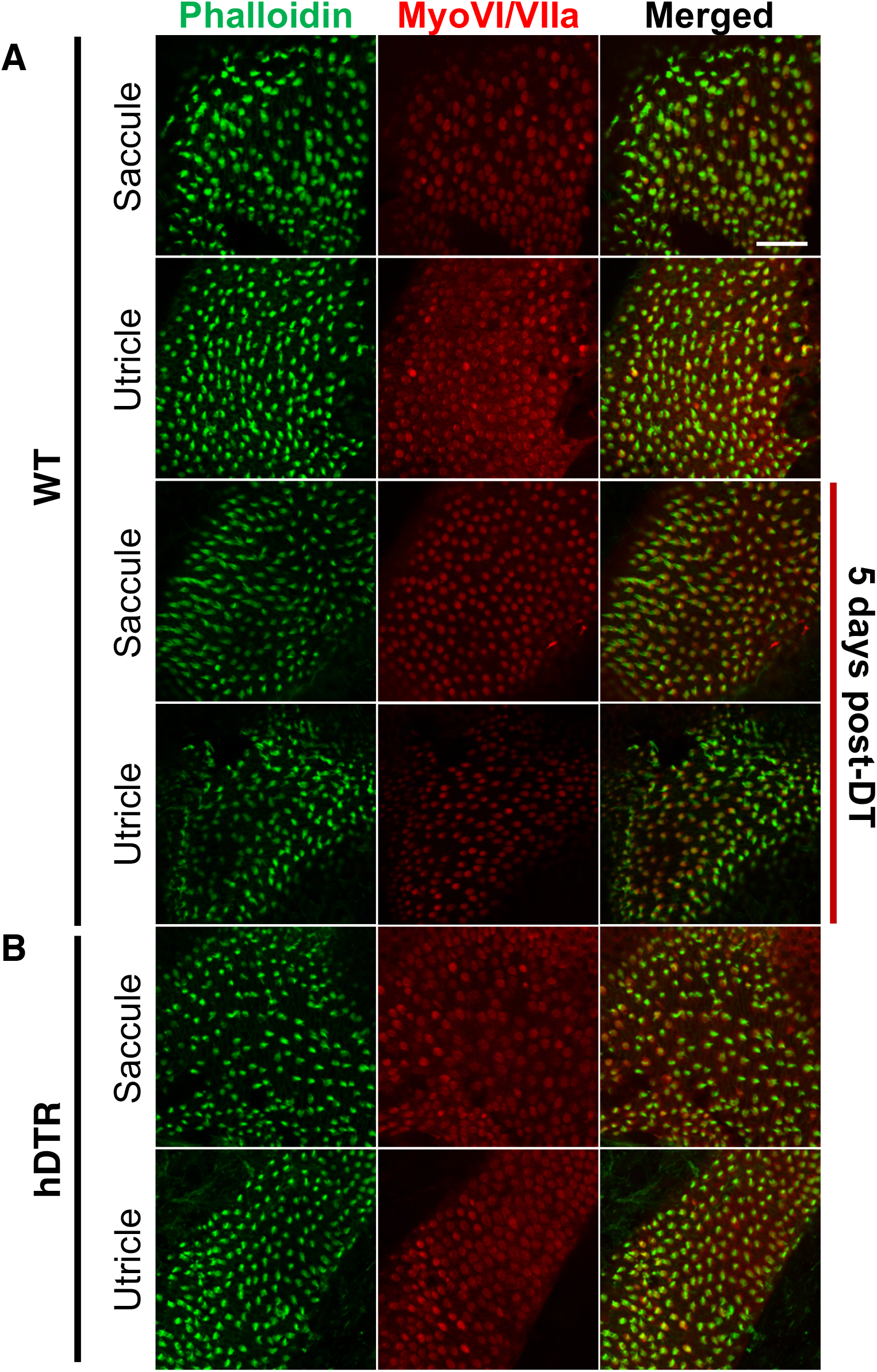
Tg(*myo6b*:hDTR) sensory epithelia. **(A)** Untreated and treated saccule and utricle from wild-type fish (WT). Saccule and utricle were isolated from treated wild-type fish 5 days post-DT. **(B)** Untreated saccule and utricle from Tg(*myo6b*:hDTR) fish (hDTR). Close examination of hair cells with phalloidin (green channel) and hair cell bodies with anti-myosin VI/VIIa (red channel). Scale bar 20 μm.

